# LevioSAM: Fast lift-over of alternate reference alignments

**DOI:** 10.1101/2021.02.05.429867

**Authors:** Taher Mun, Nae-Chyun Chen, Ben Langmead

## Abstract

**Motivation:** As more population genetics datasets and population-specific references become available, the task of translating (“lifting”) read alignments from one reference coordinate system to another is becoming more common. Existing tools generally require a chain file, whereas VCF files are the more common way to represent variation. Existing tools also do not make effective use of threads, creating a post-alignment bottleneck.

**Results:** LevioSAM is a tool for lifting SAM/BAM alignments from one reference to another using a VCF file containing population variants. LevioSAM uses succinct data structures and scales efficiently to many threads. When run downstream of a read aligner, levioSAM completes in less than 13% the time required by an aligner when both are run with 16 threads.

**Availability:** https://github.com/alshai/levioSAM

## 1 Introduction

Most analyses of sequencing datasets start by aligning the reads to a linear reference genome. But using a single linear reference leads to reference bias, a tendency to produce incorrect alignments or to miss alignments for reads containing non-reference alleles. Many methods have been proposed for reducing bias by incorporating alternate alleles into the reference (Garrison *et al.*, 2018). Some of these methods focus on aligning reads to one or more linear references containing non-reference alleles, e.g. major allele (Dewey *et al.*, 2011), or individual-specific references (Rozowsky *et al.*, 2011).

Such variant-aware methods often end with a step that translates alignments from the variant-augmented coordinate system to a standard reference. This can affect the alignment’s offset with respect to the chromosome, its CIGAR string (reflecting insertion/deletion differences), etc.

This “lift-over” problem has been addressed in prior tools including UCSC liftOver (Fujita *et al.*, 2010), and CrossMap (Zhao *et al.*, 2014). To our knowledge, only CrossMap supports lifting alignments in the SAM/BAM format (Li *et al.*, 2009). These tools also require a “chain file” as input, whereas allele information for building alternate references is usually available as VCF files (Danecek *et al.*, 2011) provided by large consortia for cataloguing variation (1000 Genomes Project Consortium *et al.*, 2015; Karczewski *et al.*, 2020).

We describe levioSAM, a new method and software tool for lifting alignments from an alternate to a target reference genome. LevioSAM uses succinct data structures to store and query the pattern of gaps in the alignment between the references. LeioSAM supports the most common file formats: SAM and BAM for alignments and VCF for genetic variants. When using multiple threads, levioSAM is faster than CrossMap but gives identical mapping results.

## 2 Approach

LevioSAM uses succinct data structures from the SDSL library (Gog *et al.*, 2014) to represent indel differences between the source and target references (Figure 1).

**Fig. 1.**
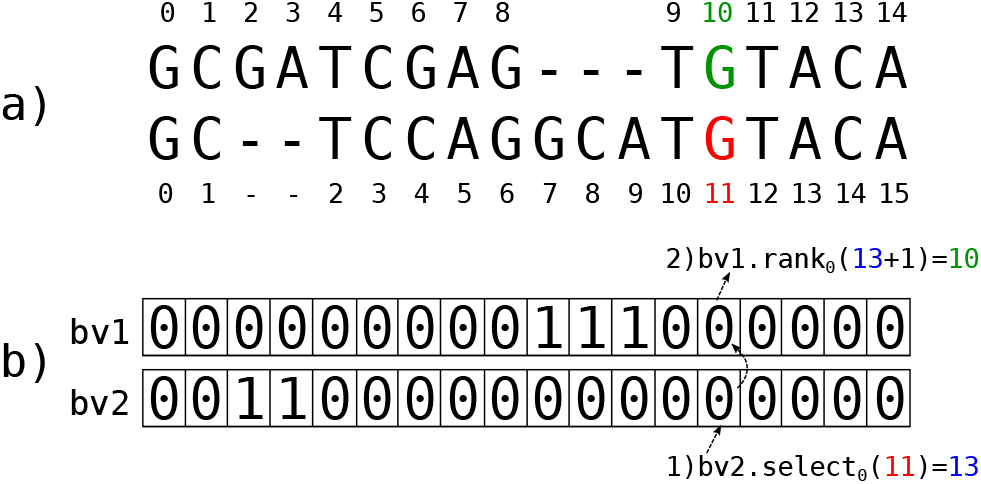
a) Example of a pairwise alignment between two sequences, with offsets labeled. The red position in the bottom sequence corresponds to the green position in the top sequence. b) visual representation of how levioSAM lifts over the red position in the bottom sequence to the top sequence. The pairwise alignment is represented by two bitvectors with 1s corresponding to gaps. (1) A select_0_ query is performed on the red position w.r.t `bv2.` (2) This value is used as input to a rank_0_ query w.r.t `bv1` to obtain the corresponding position on the top sequence.

The structures can stored on disk so they can be reused without recalculating differences. LevioSAM supports standard BAM/SAM input files. LevioSAM’s translation affects several fields of the SAM: e.g. its offset, tag information, CIGAR string, and paired-end information. VCF and BAM file parsing is supported by htslib (Li *et al.*, 2009). Multithreading and piped input/output are supported, allowing levioSAM to fit into existing workflows without creating a new computational bottleneck.

## 3 Results

We evaluated the computational efficiency of levioSAM when lifting human alignments from a major-allele version of GRCh38 to the standard GRCh38 reference. We built the major-allele reference based on the 1000 Genomes Project callset (Lowy-Gallego *et al.*, 2019), giving a linear reference with 1,746,180 single-nucleotide variants and 252,781 indels with respect to GRCh38. We randomly sampled 10 million 150-bp Illumina reads from ERR3239334, a whole-genome sequencing dataset derived from the NA12878 individual. We used Bowtie 2 (Langmead and Salzberg, 2012) with default parameters to align sequencing reads to the major-allele GRCh38 reference.

We found that levioSAM used only a fraction of the computational resources used by Bowtie 2 during alignment (Figure 2a). The mean CPU time for levioSAM is 12.0% compared to Bowtie 2 (83.6 vs. 699.3 ms per read) for single-end reads, and 12.9% (87.0 vs. 673.8 ms per read) for paired-end reads. LevioSAM used less than 13 MB memory in both cases, a fraction of Bowtie 2’s 3GB footprint. LevioSAM scaled efficiently with added threads (Figure 2b). With 32 threads, levioSAM completed 25.0 times faster than when using 1 thread and when lifting unpaired alignments, and 23.9 times faster for paired-end alignments. LevioSAM’s peak memory footprint was 54% higher for 32 threads compared to a single thread, but peak memory never exceeded 16MB.

**Fig. 2.**
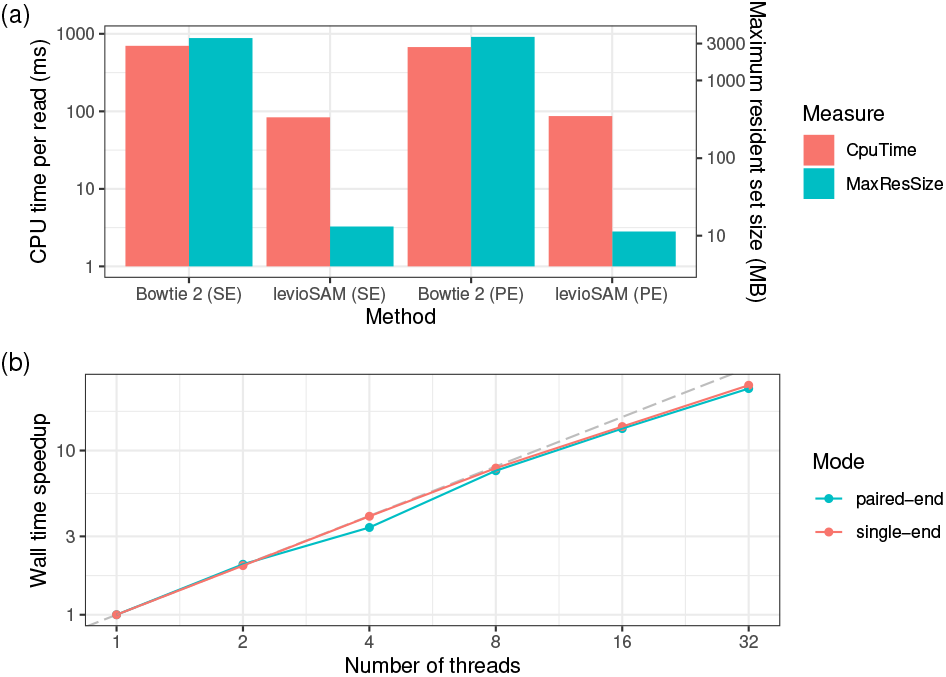
(a) Runtime of alignment and levioSAM using single-end (SE) and paired-end (PE) modes. Ten million 150-bp reads are processed using 16 threads. (b) Wall time speedup when using multiple threads. With one thread (1x), levioSAM uses 74.3 ms for a single-end read and 79.0 ms for a paired-end read segment. The gray dashed line represents the perfect scenario.

We then compared the run-time and memory of levioSAM to that of CrossMap, which also supports BAM/SAM lift-over, using the same set of reads. CrossMap requires a “chain” file rather than a VCF, so we first converted the major-allele VCF to a chain file. LevioSAM and CrossMap take a similar amount of time using a single thread (896 and 931 seconds respectively), but CrossMap lacks support for multithreading, making it a likely bottleneck. The tools produced identical lift-over coordinates for each alignment, though there were some discrepancies in the MAPQ (0.11% alignments differed) and CIGAR strings (0.12% of alignments differed) - though every CIGAR string was still valid. The MAPQ discrepancies tended to occur in discordant alignments. Overall, levioSAM’s memory efficiency, speed, and thread scaling allow it to fit in a typical read alignment workflow without creating a new bottleneck downstream of read alignment. These experiments were performed on a computer with a 2.2 Ghz Intel Xeon CPU (E5-2650 v4) and 515GB memory.

## 4 Discussion

LevioSAM is an alignment lifting tool that works with SAM/BAM and VCF files and makes it easier to implement efficient workflows for read alignment to one or more customized reference genomes. LevioSAM is based on succinct data structures and scales well with multiple threads, making it computationally efficient to be integrated in alignment pipelines. We believe this tool will further benefit methods based on augment reference genomes to reduce reference bias (Chen *et al.*, 2020). While levioSAM currently assumes that differences between the source and target references consist only of mismatches and indels, it will be important to support larger differences such as inversions or rearrangements in the future. While this is not possible in general, we expect that many cases can be handled by e.g. splitting reads into segments during the lifting process.

## Acknowledgements

Part of this research project was conducted using computational resources at the Maryland Advanced Research Computing Center (MARCC). The sequencing data we used for experiments were generated at the New York Genome Center with funds provided by NHGRI Grant 3UM1HG008901-03S1. The cell lines/DNA sample was obtained from the NIGMS Human Genetic Cell Repository at the Coriell Institute for Medical Research.

## References

1000 Genomes Project Consortium et al. (2015). A global reference for human genetic variation. Nature, 526(7571), 68.

Chen, N.-C., Solomon, B., Mun, T., Iyer, S., and Langmead, B. (2020). Reducing reference bias using multiple population reference genomes. BioRxiv.

Danecek, P., Auton, A., Abecasis, G., Albers, C. A., Banks, E., DePristo, M. A., Handsaker, R. E., Lunter, G., Marth, G. T., Sherry, S. T., et al. (2011). The variant call format and vcftools. Bioinformatics, 27(15), 2156–2158.

Dewey, F. E., Chen, R., Cordero, S. P., Ormond, K. E., Caleshu, C., Karczewski, K. J., Whirl-Carrillo, M., Wheeler, M. T., Dudley, J. T., Byrnes, J. K., et al. (2011). Phased whole-genome genetic risk in a family quartet using a major allele reference sequence. PLoS Genet, 7(9), e1002280.

Fujita, P. A., Rhead, B., Zweig, A. S., Hinrichs, A. S., Karolchik, D., Cline, M. S., Goldman, M., Barber, G. P., Clawson, H., Coelho, A., et al. (2010). The ucsc genome browser database: update 2011. Nucleic acids research, 39(suppl_1), D876–D882.

Garrison, E., Sirén, J., Novak, A. M., Hickey, G., Eizenga, J. M., Dawson, E. T., Jones, W., Garg, S., Markello, C., Lin, M. F., et al. (2018). Variation graph toolkit improves read mapping by representing genetic variation in the reference. Nature biotechnology.

Gog, S., Beller, T., Moffat, A., and Petri, M. (2014). From theory to practice: Plug and play with succinct data structures. In 13th International Symposium on Experimental Algorithms, (SEA 2014), pages 326–337.

Karczewski, K. J., Francioli, L. C., Tiao, G., Cummings, B. B., Alföldi, J., Wang, Q., Collins, R. L., Laricchia, K. M., Ganna, A., Birnbaum, D. P., et al. (2020). The mutational constraint spectrum quantified from variation in 141,456 humans. Nature, 581(7809), 434–443.

Langmead, B. and Salzberg, S. L. (2012). Fast gapped-read alignment with bowtie 2. Nature methods, 9(4), 357.

Li, H., Handsaker, B., Wysoker, A., Fennell, T., Ruan, J., Homer, N., Marth, G., Abecasis, G., and Durbin, R. (2009). The sequence alignment/map format and samtools. Bioinformatics, 25(16), 2078–2079.

Lowy-Gallego, E., Fairley, S., Zheng-Bradley, X., Ruffier, M., Clarke, L., Flicek, P., Consortium,. G. P., et al. (2019). Variant calling on the grch38 assembly with the data from phase three of the 1000 genomes project. Wellcome Open Research, 4.

Rozowsky, J., Abyzov, A., Wang, J., Alves, P., Raha, D., Harmanci, A., Leng, J., Bjornson, R., Kong, Y., Kitabayashi, N., Bhardwaj, N., Rubin, M., Snyder, M., and Gerstein, M. (2011). AlleleSeq: analysis of allele-specific expression and binding in a network framework. Mol. Syst. Biol., 7, 522.

Zhao, H., Sun, Z., Wang, J., Huang, H., Kocher, J.-P., and Wang, L. (2014). Crossmap: a versatile tool for coordinate conversion between genome assemblies. Bioinformatics, 30(7), 1006–1007.

